# Evaluation depth reshapes method rankings in proteomic co-abundance network inference

**DOI:** 10.64898/2026.11.11.765733

**Authors:** Shahram Aliyari, Nazanin Bagherlou, Zahra Salehi

## Abstract

Network inference from quantitative proteomics is usually benchmarked at a single, arbitrarily chosen number of top-ranked edges, so the influence of evaluation depth has not, to our knowledge, been systematically tested. We compared five strategies (random forest with bootstrap selection-frequency filtering, WGCNA, GENIE3, Spearman correlation and context likelihood of relatedness) across seven proteomic datasets from four quantification platforms, using harmonised preprocessing and Precision@K, the fraction of a method’s K top-ranked protein pairs present in a pooled reference of curated protein associations (STRING, BioGRID and CORUM). Method rankings changed with depth. In all six datasets that allow an equal-depth comparison, the leading method at K = 50 differed from the leader at K = 2,000, and 18 of 24 pairwise comparisons involving the filtered random forest reversed somewhere along the depth range, although seven of these reversals rested on differences of at most 0.02 in precision and each network was inferred only once. No method dominated. Relative to the same random forest without filtering, the selection-frequency filter was associated with small differences (+0.006 to +0.036 in Precision@500 in six of six equal-depth datasets; mixed at shallower depths) that cannot be separated from its 20- to 50-fold greater number of fitted trees. Prospective testing on independent datasets gave one qualified replication, one non-replication and one non-evaluable dataset. We recommend reporting precision trajectories across depths rather than single-threshold comparisons.

## Introduction

Proteins function within complex biological networks, where coordinated interactions and regulatory mechanisms are essential for maintaining cellular processes. Physical complexes, shared regulatory control and co-dependent stabilisation or degradation leave a statistical signature in quantitative proteomic data: proteins that participate in the same process tend to co-vary in abundance across samples. Networks built from these co-abundance patterns are used to prioritise functional relationships, identify hub proteins, delineate modules and generate hypotheses for targeted follow-up, particularly in cancer proteomics, where regulatory wiring is only partially known [1]. The value of such a network depends on how faithfully the inference method recovers real relationships from noisy, indirect evidence. The choice and evaluation of the method is therefore a central methodological question, not an incidental one. Inferring networks from proteomic data is more challenging than the term “co-expression network” might suggest.

Proteomic cohorts are small relative to the number of quantified proteins (typically tens to a few hundred samples against thousands of proteins), a regime in which spurious associations are common and true ones are easily missed. Missingness is pervasive and platform-specific: label-free quantification (LFQ) commonly encodes non-quantified proteins as literal zeros, whereas isobaric labelling and affinity-panel platforms have different missingness and dynamic-range characteristics. Acquisition technologies differ in noise structure and identifier system. The reference against which an inferred network can be judged is also incomplete: curated interaction databases are incomplete and non-randomly sampled [2], so agreement with a database indicates plausibility, not truth.

Several families of computational strategy address this problem. Pairwise correlation (Pearson or, more robustly, Spearman) ranks protein pairs by the strength of their monotonic co-abundance; it is fast and transparent but treats each pair in isolation. Information-theoretic methods such as the Context Likelihood of Relatedness (CLR) rank pairs by mutual information corrected against each node’s background distribution [3]. Weighted correlation network analysis (WGCNA) builds a soft-thresholded adjacency over the whole protein set and yields interpretable modules [4, 5]. Tree-ensemble methods built on the random forest [6] recast inference as one regression per target protein and rank predictors by importance; GENIE3 (Gene Network Inference with Ensemble of Trees) is the best-established example [7], with extensions that incorporate prior data [8], time series [9] and gradient boosting for scale [10].

Community benchmarks have shown that no single method dominates across datasets and that aggregating methods is more robust than trusting one [11]. The same conclusion has been reached for proteomic network inference, where a multi-method comparison across eleven cancer types found that method rankings varied by cancer type [12]. Both studies, like most benchmarks in the field, summarise each method’s performance in a single number, either at one evaluation threshold (a fixed network size, number of top-ranked predictions or low-recall window) or as an area under the precision–recall or receiver-operating-characteristic curve, and rank methods by it. Implicit in this practice is the assumption that one summary ranking holds at every depth of the ranked list. To our knowledge, this assumption has not been tested directly.

Different method families concentrate confidence in their top predictions in different ways, so it is plausible that rankings change as the number of evaluated predictions grows. If so, two otherwise identical benchmarks could name different winners simply because they chose different depths.

A parallel line of work wraps a single method in a resampling-based filter that keeps only predictions that recur across resamples. Stability selection retains only those variables or edges that recur across random subsamples above a frequency threshold and, under its assumptions (subsamples of half the samples drawn without replacement), bounds the expected number of false selections [13]. It has already been combined with per-target regression for network inference: Trustful Inference of Gene REgulation using Stability Selection (TIGRESS) applies it to a sparse-regression formulation and was a top performer in the DREAM5 challenge [14], and bootstrap edge-selection frequencies have been used to filter Gaussian graphical models [15]. What remains unexamined is whether a stability-selection wrapper changes a method’s behaviour across evaluation depths, for example by sharpening only the very top of the ranked list or by extending performance deeper, and whether any such effect is consistent across datasets.

Here we address this gap directly. We do not propose a new algorithm, and we do not claim that stability-selection wrapping of a per-target method is new. Our contribution is threefold. First, we benchmark five established strategies on seven proteomic datasets that span cancer and non-cancer biology and four quantification platforms, evaluating every method across a fixed grid of depths rather than at a single threshold, and we show how method rankings change along that grid. Second, we compare random forest with and without a bootstrap selection-frequency filter, with a matched control for candidate-pool depth, and we report that the two arms differ in computational effort. Third, we test the resulting depth-dependent hypothesis prospectively on independent datasets that were selected, and the hypothesis recorded, before any method was run on them.

This study aims to examine how evaluation depth influences the comparative performance and ranking of proteomic network-inference methods across heterogeneous datasets, and whether bootstrap selection-frequency filtering alters the depth-dependent behaviour of random-forest-based inference. By integrating retrospective benchmarking with prospective evaluation, we aim to provide a more comprehensive assessment of network-inference methods beyond a single evaluation threshold.

## Results

### A harmonised benchmark across heterogeneous proteomic datasets

We assembled eight proteomic datasets (Table 1). Five formed an initial, retrospective panel: Clinical Proteomic Tumor Analysis Consortium (CPTAC), BRCA (Breast Invasive Carcinoma), lung adenocarcinoma (LUAD), ovarian cancer (OV) and COAD (Colon Adenocarcinoma), all measured by tandem mass tag (TMT) mass spectrometry [16], and PXD005942, an LFQ pan-cancer cell-line dataset from the PRoteomics IDEntifications Database (PRIDE) repository [17]. Three further datasets were chosen afterwards for technological, biological and identifier-system diversity and were frozen before any method was run on them: PXD059074 (pancreatic adenocarcinoma, data-independent acquisition (DIA) mass spectrometry), NAFLD_SomaScan (Non-Alcoholic Fatty Liver Disease, aptamer-based SomaScan affinity proteomics [18]) and PXD018996 (kidney disease, urine proteome, LFQ). The prospective panel therefore added two quantification platforms (DIA mass spectrometry and aptamer affinity), one non-cancer context and two further identifier systems (UniProt accession and Slow Off-rate Modified Aptamer (SOMAmer) identifier) to the initial panel’s Ensembl and gene-symbol identifiers.

**Table 1.** Dataset characteristics.

| Dataset | Biological context | Platform | Samples (n) | Features (n) | Raw missingness | Identifier | Panel | Status |
|---|---|---|---|---|---|---|---|---|
| CPTAC_BRCA | Breast cancer | TMT | 122 | 3,347 | 17.7% | Ensembl gene ID | Retrospective | Evaluable |
| PXD005942 | Pan-cancer cell lines | LFQ | 58 | 271 | 70.2% | Gene symbol | Retrospective | Evaluable; excluded from cross-method summaries |
| CPTAC_LUAD | Lung adenocarcinoma | TMT | 109 | 3,243 | 7.6% | Gene symbol | Retrospective | Evaluable |
| CPTAC_OV | Ovarian cancer | TMT | 82 | 3,016 | 9.0% | Gene symbol | Retrospective | Evaluable |
| CPTAC_COAD | Colon adenocarcinoma | TMT | 97 | 2,035 | 19.3% | Gene symbol | Retrospective | Evaluable |
| PXD059074 | Pancreatic adenocarcinoma | DIA | 174 | 1,435 | 25.2% | UniProt accession (mapped) | Prospective | Evaluable |
| NAFLD_SomaScan | NAFLD (non-cancer) | SomaScan (affinity) | 189 | 2,149 | 0.0% | SOMAmer ID (mapped) | Prospective | Evaluable |
| PXD018996 | Kidney disease (urine) | LFQ | 206* | 1,721* | 79.5% | Gene symbol | Prospective | **Non-evaluable** |

Sample and feature counts for evaluable datasets are after preprocessing (missingness filtering, outlier removal and feature selection). \*PXD018996 counts are for the canonical matrix before exclusion (not benchmarked). Raw missingness is the fraction of missing cells in each dataset’s raw matrix under its own convention (explicit NA, or literal zero for the MaxQuant LFQ datasets PXD005942 and PXD018996). PXD005942’s zero-coded missingness is handled by an explicit zero-as-missing correction that reduced its features from 1,577 to 388 before feature selection (271 final); see Methods. Per-dataset filter parameters are in Supplementary Table 6.

Every evaluable dataset passed through one frozen preprocessing procedure (Fig. 1; Methods) and the resulting single matrix was supplied unchanged to all five methods: random forest with bootstrap selection-frequency filtering (RF+stability; Methods), WGCNA, GENIE3, Spearman correlation and CLR. Differences between methods therefore reflect the inference procedure, not preprocessing. Each method’s edges were scored as one candidate association per unique protein pair, so that all five methods predict the same object as the reference databases encode (Methods, “Edge representation”).

**Figure 1.**
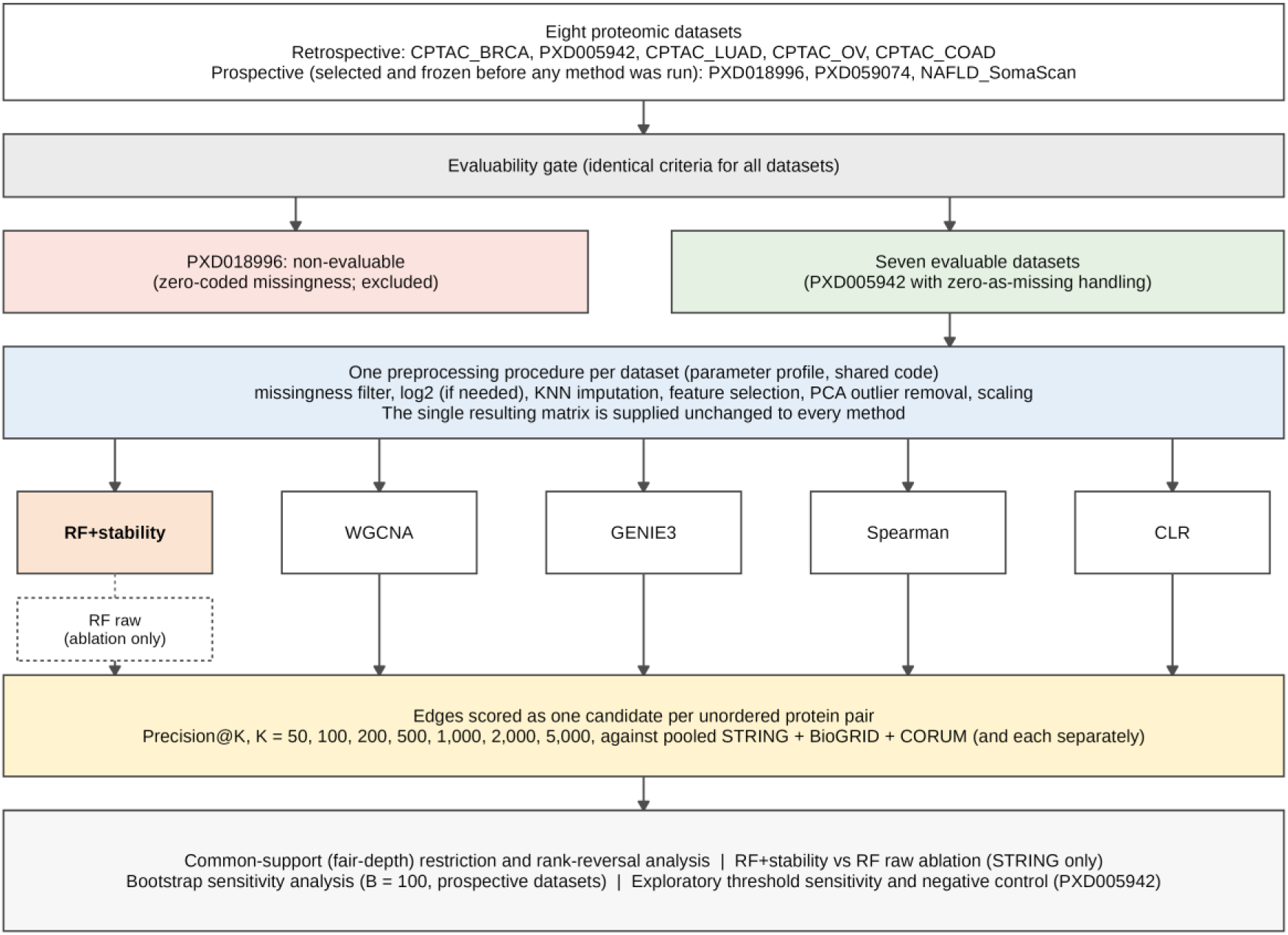
Study design. Eight proteomic datasets passed through one evaluability gate. PXD018996 was excluded as non-evaluable (zero-coded missingness incompatible with the missingness filter), leaving seven evaluable datasets; PXD005942 was analysed with explicit zero-as-missing handling (Methods). Each evaluable matrix underwent the same preprocessing, and the single resulting matrix was supplied unchanged to all five methods: random forest with bootstrap selection-frequency filtering (RF+stability), WGCNA, GENIE3, Spearman correlation and CLR. RF raw, the same random forest without filtering, is used only in the ablation. Edges were scored as one candidate per unordered protein pair by Precision@K against a pooled STRING, BioGRID and CORUM reference (and each database separately) over a fixed K grid, followed by the common-support rank-reversal analysis, the ablation, a bootstrap sensitivity analysis (B = 100) and exploratory threshold and negative-control analyses. Schematic; no plotted data. KNN, k-nearest neighbour; PCA, principal component analysis; RF, random forest.

Two datasets required specific handling, and we report both in full. PXD018996 was classified as non-evaluable: its LFQ data encode missing values as zeros, which the frozen missingness filter does not detect, and 64.6% of its 206 samples individually exceeded 80% missingness, so that 73.8% of the cells in its matrix would have been imputed. A network built from such a matrix would mainly reflect the imputation algorithm. We excluded the dataset rather than rescue it with a threshold not used elsewhere (Supplementary Table 10). Missingness patterns, filtering thresholds and imputed fractions of all eight datasets are summarised in Supplementary Table 12.

PXD005942 carries the same zero-coding. A zero-as-missing correction reduced its evaluable features from 1,577 to 388 before feature selection (271 final) and left its five inferred networks so different in size that no depth on the standard grid permits an equal-depth comparison (see “Fair depth is limited by network size” below). PXD005942 is therefore shown throughout but excluded from every cross-method summary count. Six datasets remain for the cross-method analyses.

### No method leads at shallow depth

At the shallowest depth (K = 50), the leading method differed among the six datasets: Spearman correlation led in CPTAC_BRCA and CPTAC_LUAD, WGCNA in CPTAC_OV and PXD059074, and CLR in CPTAC_COAD and NAFLD_SomaScan (Fig. 2; Supplementary Table 1). RF+stability led in none. Its rank at K = 50 ranged from 2nd (CPTAC_OV) to 5th (CPTAC_COAD, PXD059074), with 3rd in CPTAC_LUAD and NAFLD_SomaScan and 4th in CPTAC_BRCA. This heterogeneity at a single shallow depth is consistent with earlier multi-dataset benchmarks [12, 19]. We report it as the starting point for the depth analysis and because it is precisely the observation that a single-threshold benchmark at low K would present as its conclusion. Leads at K = 50 were narrow: in four of the six datasets (CPTAC_BRCA, CPTAC_LUAD, CPTAC_OV and PXD059074) the leader was ahead of the next method by a single reference-supported pair (Supplementary Table 1).

**Figure 2.**
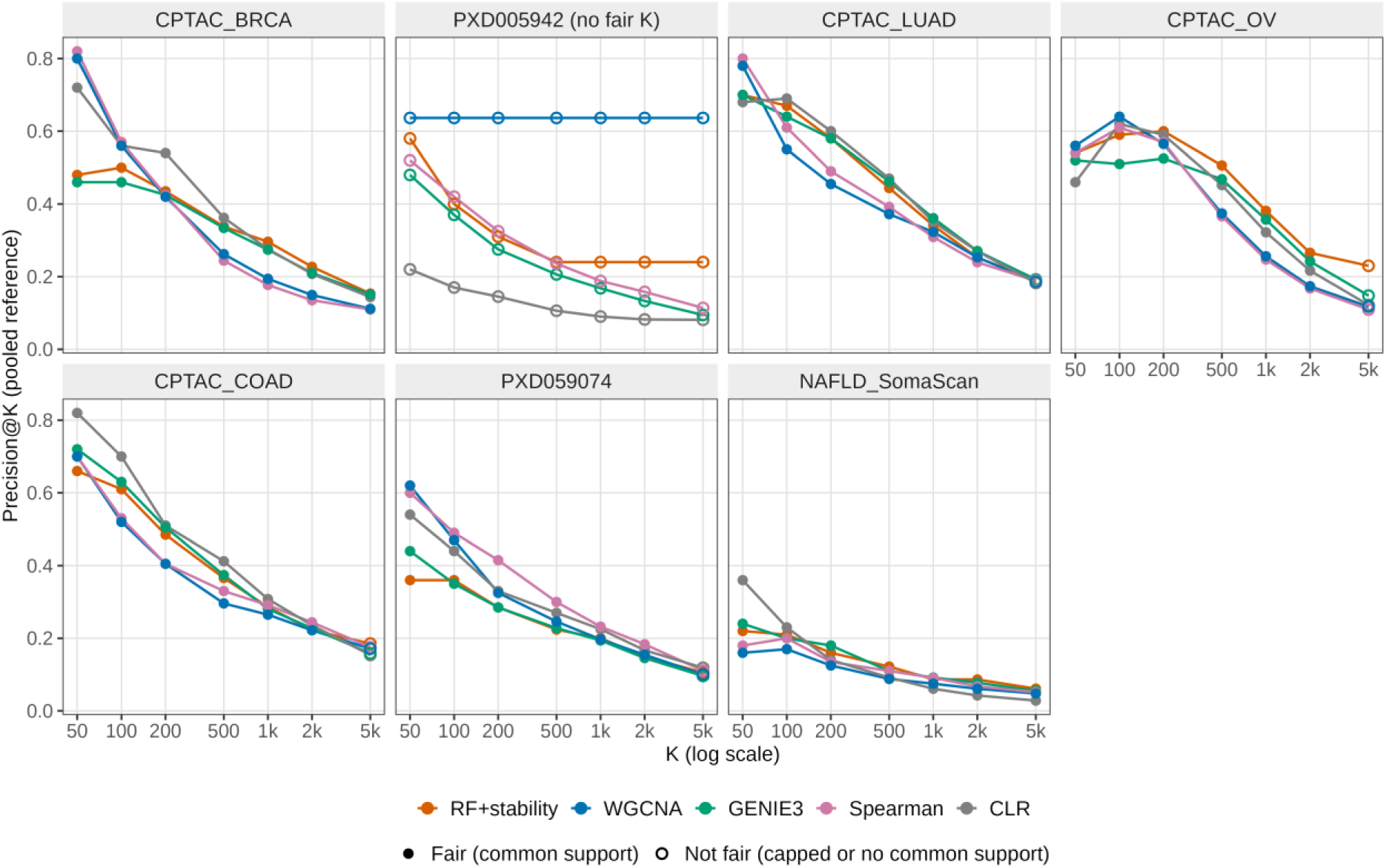
Precision@K trajectories show that no method wins at every depth. Precision@K (y-axis; fraction of a method’s top-K edges, ranked by native score, present in the pooled reference) against K (x-axis, log scale; K = 50, 100, 200, 500, 1,000, 2,000, 5,000) for the five methods (colour), one panel per dataset (CPTAC_BRCA, PXD005942, CPTAC_LUAD, CPTAC_OV, CPTAC_COAD, PXD059074, NAFLD_SomaScan). Filled markers denote fair (common-support) depths, K ≤ K_common_max; open markers denote depths that are not fair for that dataset because at least one method’s network has fewer than K pairs. One run per method per dataset (GENIE3, and the RF+stability bootstrap in five datasets, were not run under a controlled seed); point estimates, no error bars. PXD018996 is not shown (non-evaluable). PXD005942 is shown for transparency but is excluded from the cross-method summaries (Methods, common-support correction). K, number of top-ranked edges; RF, random forest.

### Evaluation depth reshapes method rankings

Extending the comparison across the fixed grid (K = 50, 100, 200, 500, 1,000, 2,000 and 5,000, restricted to each dataset’s common-support range; see below) showed that rankings are not fixed (Figs. 2 and 5a–c). In every one of the six datasets, the leading method at K = 50 was different from the leading method at K = 2,000 (Table 2). K = 2,000 is a fair depth in all six datasets because RF+stability’s network, which sets the fair range, exceeds 2,000 pairs at the primary selection-frequency threshold (see below). RF+stability reached rank 1 of 5 at some tested depth in three datasets (CPTAC_BRCA at K = 1,000, CPTAC_OV at K = 200 and NAFLD_SomaScan at K = 500) and in the remaining three (CPTAC_LUAD, CPTAC_COAD and PXD059074) never did. Trajectories were not always monotonic. In NAFLD_SomaScan the RF+stability rank across the seven depths was 3, 2, 2, 1, 3, 1, 1, so the method reached rank 1, lost it and regained it. Once RF+stability reached rank 1 in CPTAC_BRCA and CPTAC_OV, it kept it through the largest fair depth.

**Table 2.** Leading method by evaluation depth and prospective outcome.

| Dataset | Leader K=50 | Leader K=500 | Leader K=1,000 | Leader K=2,000 | First K at which RF+stability is rank 1 | Prospective outcome |
|---|---|---|---|---|---|---|
| CPTAC_BRCA | Spearman | CLR | RF+stability | RF+stability | 1,000 | Hypothesis-generating |
| PXD005942† | WGCNA | WGCNA | WGCNA | WGCNA | **Never (rank 2–3)** | Excluded (no fair K) |
| CPTAC_LUAD | Spearman | CLR | GENIE3 | GENIE3 | **Never** | Hypothesis-generating |
| CPTAC_OV | WGCNA | RF+stability | RF+stability | RF+stability | 200 | Hypothesis-generating |
| CPTAC_COAD | CLR | CLR | CLR | Spearman | **Never** | Hypothesis-generating |
| PXD059074 | WGCNA | Spearman | Spearman | Spearman | **Never** | **Non-replication** |
| NAFLD_SomaScan | CLR | RF+stability | GENIE3 | RF+stability | 500 (rank lost at K=1,000) | Qualified replication |
| PXD018996 | – | – | – | – | – | **Non-evaluable** |

“Leader” is the method with the highest Precision@K (pooled reference) at that depth; single run, no ties at these depths. Headline: RF+stability reaches rank 1 at some fair (equal-depth) K in 3 of 6 cross-method datasets (CPTAC_BRCA, CPTAC_OV, NAFLD_SomaScan) and by K ≤ 500 in 2 of 6 (CPTAC_OV, NAFLD_SomaScan); in CPTAC_LUAD, CPTAC_COAD and PXD059074 it never reaches rank 1. †PXD005942 is shown for transparency only: WGCNA’s network has 22 edges (K_common_max = 22), so its Precision@K is constant at 0.636 and no standard-grid K is a fair comparison; RF+stability ranks 2nd at all depths except K = 100 and 200 (3rd). Full ranks and capped/fair flags: Supplementary Table 1.

We quantified the pattern across all 24 pairwise comparisons between RF+stability and each other method within the six datasets (four competitors × six datasets). A comparison was classified as showing no reversal (the initial order was never inverted), a transient crossing (the order inverted at some depth but reverted before the largest fair depth) or a persistent reversal (the order inverted and never reverted); depths at which the two methods had identical Precision@K were treated as carrying no ordering information. Eighteen of 24 comparisons reversed somewhere along the depth range: ten persistently and eight transiently, and six showed no reversal (Fig. 5d; Supplementary Table 2). In seven of the 18 reversals, the reversed order never exceeded a difference of 0.02 in Precision@K (one reference-supported pair at K = 50); with a single run per method, such reversals cannot be distinguished from run-to-run variation. The split between persistent and transient reversals is less robust than the total. Under the rule originally used, which counted a tie as the second method ahead, one comparison (CPTAC_LUAD against GENIE3, tied at K = 50 and 200) was transient rather than persistent (nine persistent, nine transient), and the split also depends on where the fair depth range ends (Supplementary Table 16). Transient crossings were most frequent against GENIE3 (3 of 6 datasets), the other per-target random-forest method. Persistent reversals occurred in 3 of 6 datasets each against WGCNA, Spearman and CLR, and in one against GENIE3; in seven of the nine persistent reversals against WGCNA, Spearman and CLR, RF+stability was behind at the smallest fair depth; it was tied in one comparison and ahead in another.

The depth dependence of rankings was therefore the most consistent finding, but its direction was not uniform. We do not find that RF+stability’s relative performance improves systematically with depth. Including the capped depth K = 5,000 would have altered 2 of the 24 classifications (CPTAC_LUAD and CPTAC_COAD against CLR), which is why only fair depths are used. Descriptively, RF+stability’s gains in rank arose from both its own faster accumulation of reference-supported edges and the faster precision decline of competitors, in proportions that varied by dataset (Supplementary Table 3); we did not test these as causal mechanisms.

### Fair depth is limited by network size

A nominal depth K is an equal-depth comparison only if no method’s network has fewer than K edges; otherwise that method is scored on its whole network while denser competitors are scored on a true top-K subset (K_eff = min(K, network size)). RF+stability produced the sparsest network in six of the seven datasets (2,489–6,109 unique pairs among the six cross-method datasets), so its size sets the largest fair depth and Maximum Common-Support Depth (K_common_max): 5,804 for CPTAC_BRCA, 3,995 for CPTAC_LUAD, 2,489 for CPTAC_OV, 2,900 for CPTAC_COAD, 4,260 for PXD059074 and 6,109 for NAFLD_SomaScan. K = 5,000 is therefore capped in four of the six datasets, whereas K ≤ 2,000 is fair in all of them (Supplementary Table 11). Because RF+stability’s network size is set by its selection-frequency threshold (0.60) and its number of bootstrap iterations (B), so is the fair depth range: in an exploratory analysis of stricter thresholds, no Precision@K value at a fair depth changed, but K = 2,000 remained fair in only 4, 3, 1 and 0 of the six datasets at thresholds 0.7, 0.8, 0.9 and 0.96 (Supplementary Table 16). Rank statements above refer to fair depths only.

PXD005942 is the exception. After the preprocessing correction, WGCNA’s unmodified automatic soft-power selection produced a network of only 22 edges, so its Precision@K is identical at every K (0.636) and exceeds that of RF+stability at all depths. This is an extreme instance of the capping artefact, not a demonstration of superiority at equal depth, and it is equally not evidence in favour of RF+stability. The dataset is therefore reported as a disclosed data point and not counted towards any summary (RF+stability ranked 2nd at all depths except K = 100 and 200, where it ranked 3rd).

### Bootstrap filtering and unfiltered random forest differ little

GENIE3 implements the same per-target importance principle without bootstrap resampling, and its depth profile differed markedly from that of RF+stability (for example at K = 2,000, GENIE3 ranked 1st in CPTAC_LUAD and 5th in PXD059074). Per-target random-forest importance as implemented in GENIE3 therefore does not reproduce RF+stability’s depth profile. We compared RF+stability with the same engine’s unfiltered single-fit ranking (“RF raw”) at matched K (50, 100, 250, 500 and 1,000), using a Search Tool for the Retrieval of Interacting Genes/Proteins (STRING)-only reference for both (Fig. 3). RF raw retains 500 candidate predictors per target, whereas RF+stability records 10 per bootstrap iteration. We therefore also re-scored RF raw truncated to its top 10 per target; this changed Precision@K by at most 0.001 in any compared cell, so candidate-pool depth explains essentially none of the gap (Supplementary Table 4). Differences between RF+stability and RF raw were small at every depth and consistently positive only at K ≥ 500.

**Figure 3.**
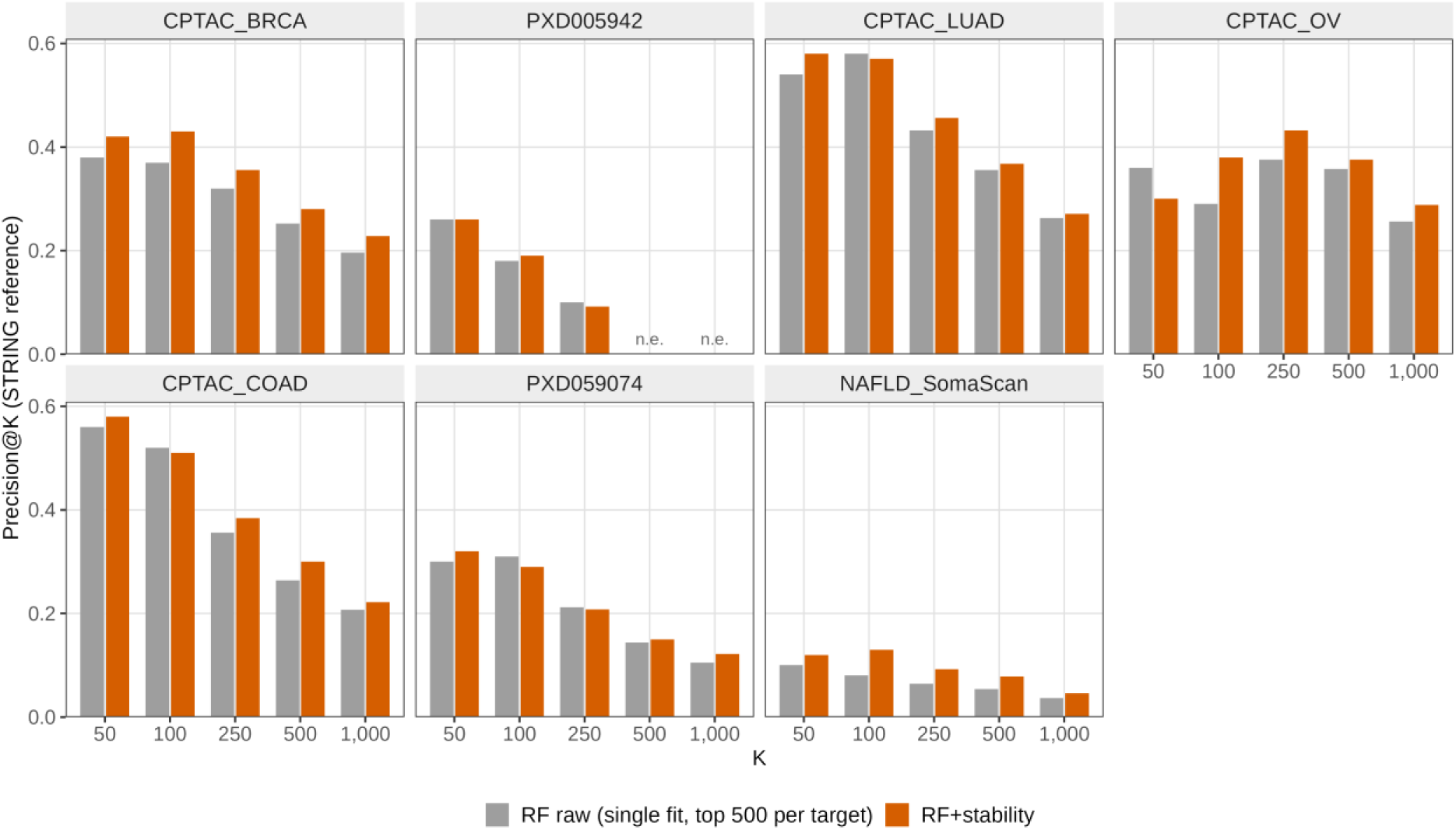
Random forest with and without bootstrap selection-frequency filtering. Precision@K (STRING-only reference) for RF+stability (B = 20, or 50 for PXD005942; selection frequency ≥ 0.60) and for the same engine’s single, untuned, unfiltered fit (RF raw; top 500 candidates per target) at K = 50, 100, 250, 500 and 1,000, one panel per evaluable dataset. n.e., not evaluated because the RF+stability network (325 pairs in PXD005942) is smaller than K. RF raw values for CPTAC_BRCA and PXD005942 are from the corrected refit (Methods). A matched control truncating RF raw to its top 10 predictors per target changed Precision@K by at most 0.001 (Supplementary Table 4). The arms differ in computational effort (20-to 50-fold more trees per target for RF+stability), so differences cannot be attributed to the filter alone. One run per arm; no error bars. RF, random forest; K, number of top-ranked pairs.

Among the six cross-method datasets, with Δ denoting the difference in Precision@K (RF+stability minus RF raw), RF+stability exceeded RF raw in five at K = 50 (all except CPTAC_OV, Δ = −0.06), in three at K = 100 (CPTAC_BRCA, CPTAC_OV and NAFLD_SomaScan; the others were lower by 0.01– 0.02), in five at K = 250 and in all six at K = 500 and K = 1,000 (Fig. 3; Supplementary Table 4). The differences at K ≥ 500 were small (Δ = +0.006 to +0.036 at K = 500 and +0.008 to +0.032 at K = 1,000). In CPTAC_BRCA, Precision@100 was 0.37 without and 0.43 with filtering (Δ = +0.06); in PXD005942 the differences were 0.000, +0.010 and −0.008 at K = 50, 100 and 250, and K = 500 and 1,000 were not compared because the filtered network has only 325 pairs. These values replace those of an earlier version, in which a configuration defect had fitted the RF raw arm of CPTAC_BRCA and PXD005942 with one candidate predictor per split (Methods). The comparison is not effort-matched: RF+stability fits 20 times (50 times for PXD005942) as many trees per target as RF raw and resamples the data, so the differences cannot be attributed to the frequency filter alone. Each value comes from a single run.

### Reference support is non-degenerate, and the choice of reference matters

Against the pooled STRING, Biological General Repository for Interaction Datasets (BioGRID) and Comprehensive Resource of Mammalian Protein Complexes (CORUM) reference (1,017,052 pairs), RF+stability had the highest support at K = 1,000 in two of six datasets (CPTAC_BRCA, 0.296; CPTAC_OV, 0.381) and ranked third in the remaining four (CPTAC_LUAD, 0.341; CPTAC_COAD, 0.286; PXD059074, 0.200; NAFLD_SomaScan, 0.087; Fig. 4). Gene-symbol mapping, applied only at the validation boundary and never inside the inference matrix, resolved 95.2– 100% of analysis-matrix features to gene symbols; unresolved placeholders were retained and cannot match the reference (Supplementary Table 8). Support fractions were lower against CORUM alone than against STRING alone or the pooled set, reflecting the size and scope of each database and not a property of any method (Supplementary Table 7). We term this “reference support” and not biological truth. Because reference coverage of the measured proteins differs between datasets, absolute Precision@K is best compared within, not between, datasets: the chance level, the fraction of all pairs of measured proteins present in the reference, was 0.9–1.5% in the six cross-method datasets and 6.7% in PXD005942, and every method exceeded it at every fair depth, by at least 2.1-fold (Supplementary Table 7c).

**Figure 4.**
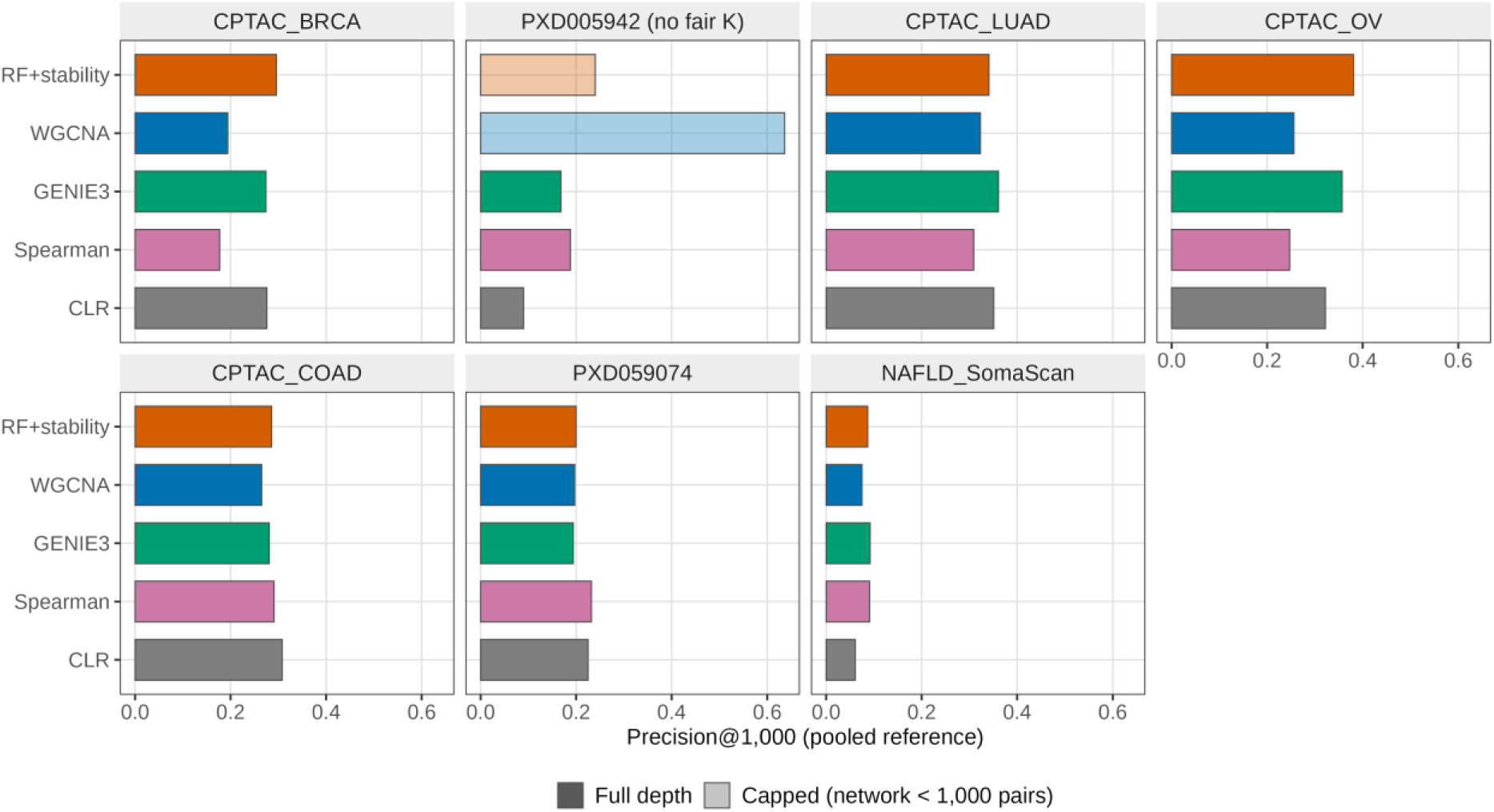
External reference support at K = 1,000 is non-degenerate for every method, and no method leads in most datasets. Precision@1,000 (x-axis; fraction of each method’s top 1,000 edges present in the pooled STRING, BioGRID and CORUM reference) for the five methods (y-axis, horizontal bars; colour by method), one panel per dataset, all seven evaluable datasets. Solid bars denote fair comparisons; faded bars denote capped comparisons in which the network has fewer than 1,000 edges (RF+stability and WGCNA in PXD005942). One run per method per dataset; point estimates. Agreement with a reference database indicates plausible, non-degenerate recovery and is not evidence of physical interaction. K, number of top-ranked edges; RF, random forest.

The choice of reference itself shifted a headline result, in the same way that depth does. CPTAC_OV reached rank 1 by K = 500 under every reference. NAFLD_SomaScan reached rank 1 at K = 500 under STRING alone or the pooled set, but not until K = 2,000 under BioGRID alone and K = 5,000 under CORUM alone (Supplementary Table 7). The reference database used for evaluation is therefore a second consequential benchmarking choice.

### Prospective evaluation gives a qualified replication, a non-replication and a non-evaluable dataset

The depth hypothesis was recorded in project documentation before the three prospective datasets were selected, and those datasets were frozen before any method was run on them (Methods). The original hypothesis was that RF+stability is not leading at low K but reaches rank 1 by K ≤ 500 in most datasets. After the edge-representation correction, the retrospective panel itself no longer showed this uniform pattern: RF+stability reached rank 1 by K ≤ 500 in only one of four retrospective datasets (CPTAC_OV), reached it later in a second (CPTAC_BRCA, K = 1,000) and never reached it in the other two. The prospective results below therefore test a weaker, less uniform pattern than originally stated.

NAFLD_SomaScan (rank 3, 2, 2, 1, 3, 1, 1 across the grid) reached rank 1 at K = 500, within the hypothesised boundary of K ≤ 500, but not monotonically. We classify this as a qualified replication of the depth-boundary criterion only. PXD059074 (rank 5, 4, 4, 5, 3, 4 from K = 50 to 2,000; K = 5,000 is outside its fair range) never reached rank 1 and did not improve monotonically; we classify it as a non-replication, not a partial one. PXD018996 could not be evaluated, which we regard as informative: a dataset chosen in good faith for diversity proved structurally incompatible with the frozen protocol, and this was disclosed instead of replaced after the fact. Because the outcome categories were written down after these results were available (Methods), the classifications are descriptive; the full rank trajectories are reported so that readers can apply their own criteria.

### Sensitivity analyses

The primary stability-selection procedure does not persist its per-iteration edge lists, so a full resampling-based uncertainty analysis of the RF+stability network was not tractable. As a disclosed sensitivity analysis we drew B = 100 sample-level bootstrap resamples of each prospective dataset and fitted every method once per resample (random forest as a single, untuned fit, architecturally equal to RF raw), using identical resamples across methods. This estimator is narrower than the primary analysis and must not be read as its uncertainty quantification. Paired differences and Holm-corrected [20] p-values are in Supplementary Table 5.

In NAFLD_SomaScan, the single-fit random forest was significantly behind CLR at K = 50 (mean difference −0.235, adjusted P = 0.040, the smallest value attainable with B = 100 after Holm correction) and K = 100 (−0.121, adjusted P = 0.040), but not at K = 500 (−0.002, adjusted P = 0.436) or K = 1,000 (+0.008, adjusted P = 0.297). The comparison with CLR is thus indistinguishable from zero at precisely the depths nearest the primary analysis’s rank-1 crossover. In PXD059074 at K = 1,000, the single fit was significantly behind WGCNA, Spearman and CLR (adjusted P = 0.040 each) and directionally, but not significantly, behind GENIE3 (adjusted P = 0.198), consistent with the absence of a rank-1 crossover in the primary analysis. We regard this analysis as supportive, not definitive. Sensitivity analyses showed that increasing the selection-frequency threshold progressively reduced the number of retained (directed) edges in all six datasets examined (Supplementary Table 13), while the distribution of stability scores varied between datasets. In addition, perturbing the number of random-forest trees from 300 to 240 produced only small changes in Precision@K across the evaluated depths (at most 0.03; Supplementary Table 13), although a single perturbation in one dataset does not establish robustness to hyperparameter choice in general.

### Selection frequency is not evidence against chance association

To calibrate what the selection frequency means, we repeated the RF+stability procedure on PXD005942 after permuting each protein independently across samples, which removes every association while preserving each protein’s distribution (20 permutations; exploratory analysis). At the primary threshold of 0.60 the procedure returned 138 ± 13 unordered pairs (range 112– 156) per permuted dataset, compared with 325 in the observed data; at thresholds of 0.80 and 0.96 it returned 16 ± 4 and 1.0 ± 1.0 pairs, compared with 114 and 41 (Supplementary Table 17). The maximum selection frequency reached 1.0 in 7 of 20 permutations. The reference overlap of these null networks was low (Precision@K 0.07–0.09 at K = 22–100, close to the chance level of 0.067 in this dataset, compared with 0.40–0.59 for the observed network). A high selection frequency therefore shows that an edge is reproducibly selected when the observed samples are resampled, not that it reflects a true association, and the 0.60 threshold provides no false-positive control. This analysis covers one dataset and one configuration and does not estimate a false-discovery rate for the observed networks.

**Figure 5.**
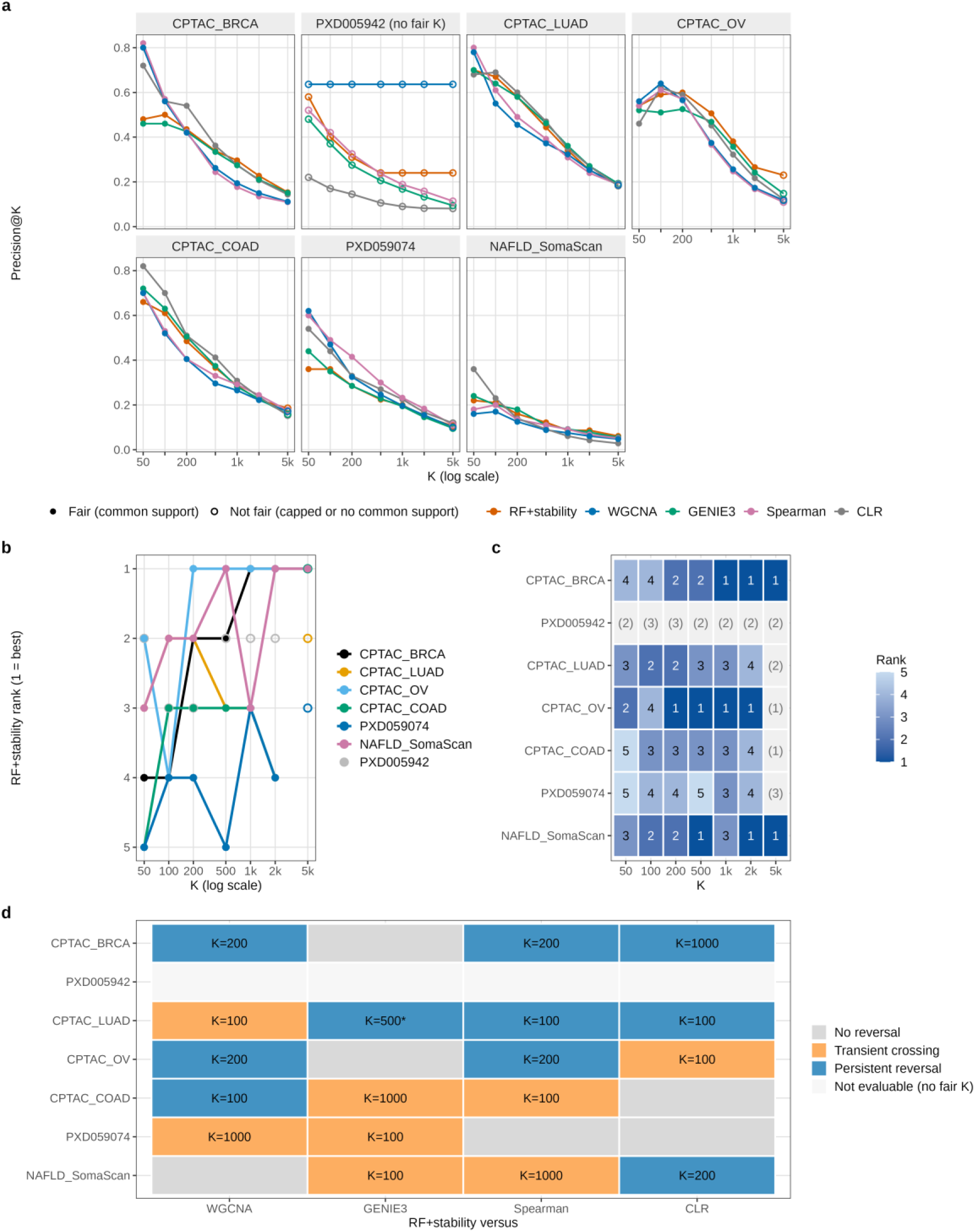
Evaluation depth reshapes the rank of RF+stability: 18 of 24 pairwise comparisons show a reversal. **a**, Precision@K trajectories for all methods and datasets (colours as in Fig. 2; open markers, depths that are not fair). **b**, Rank of RF+stability among the five methods (1 = best) against K; lines join fair depths and open markers show depths that are not fair (including all depths of PXD005942). CPTAC_LUAD, CPTAC_COAD and PXD059074 never reach rank 1 at a fair depth. **c**, Ranks in b as a heatmap; ranks at depths that are not fair are shown in parentheses on a grey background. **d**, Classification of RF+stability against each other method at fair depths, with depths at which the two methods tie treated as carrying no ordering: grey, no reversal; orange, transient crossing; blue, persistent reversal; label, first K at which the order reverses; *, classification differs under the rule originally used (a tie counted as the second method ahead), under which CPTAC_LUAD versus GENIE3 is a transient crossing (Supplementary Table 2). PXD005942 is not evaluable (no fair grid depth). One run per method per dataset; no error bars, resampling or hypothesis tests. K, number of top-ranked pairs; RF, random forest.

## Discussion

Our central finding is that the ranking of network-inference methods applied to proteomic data is not invariant to evaluation depth. Crossing precision curves are expected whenever methods concentrate their reference-supported predictions differently along their rankings; what this study adds is a measurement of how often such crossings change the conclusion of a proteomic benchmark when comparisons are restricted to equal depths, and the finding that they do so in most comparisons. A benchmark run only at K = 50 and one run only at K = 2,000 would have named different winners in every one of the six datasets that permit equal-depth comparison at the primary threshold, and 18 of 24 pairwise comparisons with RF+stability reversed somewhere along the grid. The finding does not contradict earlier benchmarks. Instead it adds an orthogonal axis to their conclusion that no method dominates across datasets[11, 12]: dataset identity and evaluation depth both determine which method appears best. We did not find prior work reporting rank reversals across a swept depth range in proteomic network inference; the standard benchmarking framework for single-cell transcriptomic inference summarises performance by the area under the precision–recall curve and early precision [21]. Our literature search was necessarily bounded, and we claim only that the pattern was not reported, not that it is absent from earlier work.

We offer several candidate explanations, none of which we tested against the others. Strong correlations in proteomic data often arise from tightly co-regulated clusters, such as members of one complex. A correlation-or mutual-information-based method could therefore concentrate much of its reference-supported yield in its first few dozen predictions. Random-forest importance aggregates over many predictors per target and may spread its signal over a longer list, which would be consistent with the shallower decline of its precision curves (Fig. 2); in a related cross-omics comparison, a random-forest method (GENIE3) produced networks of similar accuracy to correlation-and mutual-information-based methods while linking more features across data types [22]. Scores from the five methods are on incomparable numeric scales, so ties or near-ties may cluster differently at low K, even though our rank-based metric avoids a shared threshold. Bootstrap filtering could favour edges that are reproducibly selected over individually strong ones, which matters more as the list lengthens, although reproducible selection is not evidence of association (see the negative control above). Finally, reference databases are incomplete and non-randomly sampled[2], if they over-represent well-studied proteins and the strongest pairwise correlations preferentially involve such proteins, correlation-based methods may enjoy an ascertainment advantage at low K that is shared across all methods and not specific to any one. Related work has shown that apparent performance also depends on the topological scale at which networks are evaluated[23, 24], suggesting that “evaluation choice affects apparent performance” recurs along more than one axis.

The contribution of bootstrap selection-frequency filtering is small and, in our design, not separable from computational effort. Wrapping a per-target importance method in a resampling filter is an established design[13, 14, 25], and we do not present it as new.

What our ablation adds is a depth-resolved view: RF+stability exceeded unfiltered random forest in all six comparable datasets at K = 500 and K = 1,000, by 0.006 to 0.036 in Precision@K, whereas the sign of the difference varied between datasets at K ≤ 250. Because the filtered procedure fits 20 to 50 times more trees per target and resamples the data, an ensemble-size-matched comparison would be needed to attribute these differences to the filter. Users should check the contribution of such a wrapper on their own data at the depth they intend to use, not assume it from precedent.

The prospective component is, in our view, the most methodologically important, regardless of outcome. Because the hypothesis was recorded and the datasets frozen before any method was run on them, the results carry more evidential weight than a retrospective re-analysis, although the outcome categories were defined only after the results were available. They were not strongly supportive: one qualified replication, one non-replication and one dataset that proved non-evaluable. Taken together with the weakening of the retrospective pattern after correction of the edge representation, this indicates that rank-1 convergence of RF+stability at a shallow fixed K is not a reliable property of the method. We consider this an informative result and report it without softening. A retrospective benchmark would not have produced it: it would simply never have included PXD018996.

The practical implication is that method choice should be informed by the number of top-ranked relationships the downstream analysis will actually use. A researcher who will follow up only a few dozen candidate pairs should not assume that the method that leads a benchmark at K = 2,000 is best at their depth, nor the reverse. We suggest that proteomic network-inference benchmarks report full precision trajectories over a pre-specified depth grid, flag depths at which any method’s network is capped by its own size, and state the reference database used, since each of these choices can change a headline conclusion.

Our study has limitations that qualify these claims. Reference databases are incomplete and ascertainment-biased proxies, and the networks inferred here reflect statistical co-abundance, which cannot distinguish direct interaction from shared upstream regulation or confounding, including sample-level structure such as tissue of origin or tumour purity. Feature selection retained the more variable and abundant proteins (32–43% of missingness-filtered proteins in the six cross-method datasets; Supplementary Table 6), which favours well-studied, reference-rich proteins. The bootstrap analysis uses a narrower single-fit estimator and was run for the two prospective datasets only. Only two of three prospective datasets were evaluable and neither replicated the original pattern cleanly. Technology and biology strata are small (a single affinity platform and a single non-cancer context), and cancer datasets form the majority of the panel, so platform-or biology-specific statements remain descriptive.

Further limitations concern the analysis design. Random-forest settings were not uniform: PXD005942 used 500 trees, impurity importance and B = 50, and the other six datasets 300 trees, permutation importance and B = 20 (Supplementary Table 14). A 20% reduction of the tree number (300 to 240) in PXD059074 changed Precision@K by at most 0.03, but only one dataset and one direction were tested. No formal hypothesis test was applied to the rank-reversal analysis, which is descriptive. Furthermore, a rank reversal does not necessarily imply a large difference in predictive performance, because small changes in Precision@K can alter the ordering of closely performing methods. The reversal counts should therefore be interpreted as descriptive evidence of ranking instability across evaluation depths, rather than as evidence that every reversal represents a substantial performance difference. GENIE3 and the bootstrap stage of RF+stability in five datasets were run without a controlled global random seed, so exact reruns will not reproduce identical edge lists, although a seeded check confirmed that both are otherwise deterministic. A sixth method in the pipeline (a graphical-lasso approach) was excluded from this comparison as a pre-existing scope decision. The selection frequency of RF+stability is a ranking heuristic without false-positive or false-discovery control: in a permutation negative control on one dataset, a network about two-fifths the size of the observed one passed the 0.60 threshold, and null edges were retained on average even at stricter thresholds.

Because RF+stability’s network size defines the fair depth range in six of seven datasets, the largest fair depths include its lower-frequency edges, which in that control were the most compatible with chance; this affects the interpretation, not the values, of large-K comparisons and was not tested in the other datasets. The ablation compares procedures that differ in computational effort as well as in filtering. Spearman correlation was scored by its signed coefficient, so strong negative correlations were ranked last (Methods).

Future work should proceed along four lines: a larger prospective panel covering more technologies and biological contexts, to test whether platform or biology, rather than dataset idiosyncrasy, explains differences in the depth at which rank 1 is reached; a tractable way to persist and resample the primary stability-selection replicates, which would permit formal uncertainty quantification of the reversal pattern; an ensemble-size-matched ablation and negative controls in further datasets; and direct tests of the candidate mechanisms above, for example by stratifying edges by annotation depth of their endpoints or by the linearity of their co-abundance.

In summary, evaluation depth is an under-examined and consequential axis of proteomic network-inference benchmarking. RF+stability is not the best method at every depth or in every dataset, and its rank relative to other methods can rise, fall or oscillate with depth. Relative to unfiltered random forest, its selection-frequency filter was associated with small differences that we cannot separate from its greater computational effort. Single-threshold comparisons cannot distinguish a uniformly strong method from one that is strong only in the region of the ranking that the benchmark happened to examine.

## Methods

### Study design

The study combines a retrospective five-dataset initial panel (CPTAC_BRCA, PXD005942, CPTAC_LUAD, CPTAC_OV, CPTAC_COAD) with a prospective three-dataset panel (PXD018996, PXD059074, NAFLD_SomaScan), analysed under one frozen protocol. The initial panel generated the depth-dependent hypothesis. The prospective datasets were then selected for biological, technological and identifier-system diversity, and the selection was recorded before any method was run on them (time-stamped project documentation; no external registry was used). Changes made after results had been observed were limited to corrections of identified implementation defects (listed under Statistics, software and reproducibility), each applied to all affected results; in addition, the prospective outcome categories were written down after the prospective results were available (Prospective replication design). Where a parameter required judgement (for example feature-selection stringency), it was set on measured computational cost alone before biological output was inspected.

### Datasets and inclusion criteria

Datasets had to provide protein-level quantitative abundance across biologically independent samples, a resolvable identifier system and metadata sufficient to separate biological from technical replicates. CPTAC_BRCA, CPTAC_LUAD, CPTAC_OV and CPTAC_COAD (TMT) were obtained as gene-level matrices from LinkedOmics [26] and derive from the original CPTAC studies [27–30]. PXD005942 (LFQ, National Cancer Institute 60 Human Tumor Cell Line Panel (NCI-60) [31]), PXD018996 (LFQ, urine) and PXD059074 (DIA, pancreatic adenocarcinoma) were obtained from PRIDE/ProteomeXchange [17]. NAFLD_SomaScan was obtained from the supplementary data of ref. [32]. PXD018996 met all inclusion criteria at selection; its exclusion resulted from the evaluability assessment applied identically to every dataset.

### Preprocessing

Each dataset’s canonical feature-by-sample matrix was built from its native format by a source-specific adapter that performed extraction and reshaping only, and was checksummed Secure Hash Algorithm 256 (SHA-256) against a frozen manifest. Every matrix then passed through one downstream procedure applied via a dataset-specific parameter profile (not dataset-specific code): (i) missingness filtering (non-missing fraction ≥ 0.50–0.60 according to the calibrated profile); (ii) log2 transformation where the native scale was not already logarithmic (PXD059074 was recognised as log2-scale by its negative values); (iii) k-nearest-neighbour imputation with k = 10 [33]; (iv) optional joint variance-and mean-abundance-quantile feature selection, applied identically to all methods in a dataset and calibrated on compute cost alone; (v) outlier removal by principal component analysis (PCA; samples whose Euclidean distance over the first five principal-component scores exceeded the mean distance by more than 3 s.d.); and (vi) centring and unit-variance scaling. Source-study normalisation (for example CPTAC TMT ratios, SomaLogic internal normalisation) was neither undone nor repeated. Duplicate feature identifiers were removed by retaining the first occurrence, and the step was logged.

By default the missingness filter evaluates is.na() on the raw matrix before log2 transformation. This is correct for six of seven evaluable datasets, whose missing values are NA-coded or absent. MaxQuant LFQ intensities in PXD005942, however, encode “not quantified” as zero (70.2% of its raw cells; 75.4% of its 1,577 proteins exceeded 50% true missingness), which the default filter does not detect. We added an explicit zero_is_missing flag to PXD005942’s profile only, which reduced the evaluable features from 1,577 to 388 before feature selection (271 final). The same issue, more severe, underlies the non-evaluability of PXD018996: under the unmodified filter no features were removed (1,721 of 1,721 retained), 64.6% of its 206 samples individually exceeded 80% missingness, and 73.82% of cells in the resulting zero-variance-filtered matrix (1,340 features) would have required imputation. We did not apply the flag to PXD018996 and re-assess, because its missingness is far more severe than that of PXD005942 and no plausible correction would alter its classification; this is a scope decision, not an inconsistency.

### Identifier mapping

CPTAC_LUAD, CPTAC_OV, CPTAC_COAD and PXD005942 use gene symbols natively (PXD018996 after adapter-level mapping); CPTAC_BRCA uses Ensembl gene IDs; PXD059074 uses UniProt accessions (98.00% mapped, 112 retained as UNMAPPED placeholders); and NAFLD_SomaScan uses SomaLogic SOMAmer identifiers (99.28% mapped, 4,996 of 5,032). Mapping to gene symbols was applied only at the external-validation boundary and never inside the inference matrix. Ensembl IDs were mapped with org.Hs.eg.db through AnnotationDbi after stripping version suffixes, and unmapped IDs retained their original identifier so that counts remain accurate. For the 232 gene symbols targeted by more than one SOMAmer, a rule was fixed before any validation result was inspected: an edge between two SOMAmers mapping to the same gene becomes a self-loop and is excluded from validation scoring (45 pairs, reducing 6,154 collapsed pairs to 6,109), whereas an edge between SOMAmers of different genes is scored as that gene pair without cross-pair deduplication.

### Random forest with bootstrap selection-frequency filtering (RF+stability)

For each target protein, abundance was regressed on all other proteins using ranger [34] with minimum node size 5, mtry = ⌊√p⌋ (p, the number of predictors), the same fixed seed for every fit and single-threaded execution. Six datasets used 300 trees and permutation importance; PXD005942 used 500 trees and impurity importance (Supplementary Table 14). The top 500 predictors per target by importance were retained as directed candidate edges; this single, untuned fit is “RF raw”. In CPTAC_BRCA and PXD005942 an internal tuning step had been enabled for this arm; because of a configuration defect it fitted RF raw with one candidate predictor per split, so both RF raw networks were refitted without tuning using the same base settings as RF+stability. The RF+stability networks were never tuned and are unaffected.

RF+stability wraps the same untuned procedure in a bootstrap selection-frequency filter, in the spirit of stability selection [13] but without its subsampling scheme or error control. For each of B bootstrap iterations (B = 20; B = 50 for PXD005942), samples were resampled with replacement, the per-target forests were refitted and each target’s 10 highest-importance predictors were recorded. An edge’s selection frequency is the fraction of the B iterations in which it was recorded. Edges with a frequency of at least 0.60, a pipeline default fixed before any benchmark result was obtained, form the RF+stability network, ranked by selection frequency and then by mean importance over the iterations in which the edge was selected. The procedure is a ranking heuristic: the selection frequency measures how reproducibly an edge is selected when the observed samples are resampled, not the probability that it is real, and no false-positive or false-discovery control is implied. RF+stability fits B times as many trees per target as RF raw (6,000 versus 300; 25,000 versus 500 for PXD005942); its recorded wall-clock time was 1.4–3.9 h per cross-method dataset, against 6–17 min for GENIE3 and seconds for WGCNA (Supplementary Table 9).

### Edge representation for cross-method scoring

WGCNA, Spearman and CLR yield one symmetric value per protein pair. GENIE3 and RF+stability (and RF raw) are per-target regression methods that natively give up to two directed values per pair. Before any scoring, GENIE3 was collapsed to the maximum of each pair’s two importance values, and RF+stability and RF raw to the higher-ranked direction (by selection frequency, then mean importance). Every method is thus scored on one candidate association per unique unordered pair, matching the encoding of STRING, BioGRID and CORUM. This convention was applied identically to all results in this paper. It replaced an earlier scoring in which RF+stability’s two directions could occupy two ranked positions, which inflated its effective depth; that earlier scoring changed RF+stability’s rank in 21 of 42 dataset-by-depth cells and is retained only as a historical sensitivity comparison (Supplementary Table 15).

### WGCNA, GENIE3, Spearman and CLR

WGCNA[4, 5] used a signed-hybrid adjacency on the transposed matrix with soft-threshold power chosen automatically by pickSoftThreshold under the signed network type (powers 1–10 and even powers 12–20, falling back to 6), Pearson correlation with pairwise-complete observations, and a minimum adjacency of 0.01; edge weight is the soft-thresholded adjacency. GENIE3 [7] used the Bioconductor implementation (RF engine, GENIE3’s K = “sqrt” setting for the number of candidate predictors per split, 300 trees; 500 for PXD005942; no controlled random seed), symmetrised by the maximum and retaining pairs with positive importance; it has no resampling or stability filter and serves as the comparator that isolates the importance principle. Spearman correlation used pairwise-complete observations and ranked pairs by the signed coefficient, so that strong negative correlations were ranked last; this convention was used for all reported results. CLR [3] discretised each protein into 8 quantile bins, computed pairwise mutual information (MI), converted it to node-wise z-scores z_i = max(0, (MI_ij − μ_i)/σ_i), and scored each pair as √(z_i² + z_j²), where MI_ij is the MI of proteins i and j and μ_i and σ_i are the mean and standard deviation of protein i’s MI values. All methods were run single-threaded.

### Harmonised benchmarking

Native network sizes differ by orders of magnitude (from hundreds or thousands of edges for RF+stability to millions for Spearman, CLR and GENIE3), so edge counts, overlap and Jaccard indices were not used. Each method was instead scored by Precision@K at K ∈ S, where S denotes the set of values 50, 100, 200, 500, 1,000, 2,000, and 5,000, a grid that was the pipeline default before any prospective result was available (retrospective results at these depths had been seen before the depth framing was adopted). For a method with network size n, the evaluable depth is K_eff = min(K, n), and Precision@K is the fraction of its top-K_eff edges, ranked by native score, whose order-independent gene-symbol pair is in the reference set. The rank of a method at a dataset and depth is its ordinal position among the five methods (1 = best), with ties given the minimum rank.

### Common-support correction and rank-reversal analysis

A depth K is a fair, equal-depth comparison only if K ≤ K_common_max, the smallest native network size among the five methods in that dataset. We computed K_common_max from recorded network sizes and flagged every dataset × method × depth cell as capped or fair without rerunning any method. For PXD005942, K_common_max = 22 (WGCNA), below every grid value, so the dataset is excluded from all common-support analyses. In the other six datasets K_common_max is set by RF+stability and therefore depends on its selection-frequency threshold (Supplementary Table 16).

For each method pair in each dataset, the order at the smallest fair depth was compared with the order at every larger fair depth. A pair showed no reversal if the order was never inverted, a transient crossing if it inverted and reverted before the largest fair depth, and a persistent reversal if it inverted and did not revert. Depths at which the two methods had identical Precision@K were treated as carrying no ordering information and were skipped, a rule adopted after the results were available; the classification under the rule originally used (a tie counted as the second method ahead) is reported alongside (Supplementary Table 2). The 24 comparisons of RF+stability against the other four methods in the six common-support datasets are reported in the Results. Including capped depths would have changed 2 of the 24 classifications (CPTAC_LUAD and CPTAC_COAD against CLR, both at K = 5,000); only the common-support version is reported.

### External validation

The pooled reference set was built once, identically for all datasets, from STRING [35] (v12.0 physical subnetwork, combined score ≥ 700; 86,519 pairs; according to STRING’s documentation, edges in this subnetwork denote membership of a physical complex, and scores combine several evidence channels), BioGRID [36] (release 5.0.255, physical and genetic interactions, including 16,429 pairs supported only by genetic evidence and 3,988 self-pairs, which cannot affect precision) and CORUM [37] (release file downloaded on 24 March 2026; all pairs within annotated complexes, that is, co-membership rather than direct contact; 57,692 pairs), normalised to uppercase, order-independent gene-symbol pairs and pooled non-redundantly (1,017,052 pairs). BioGRID alone contained 944,068 pairs. Each source was also scored separately. Edges whose endpoints could not both be mapped were excluded from scoring only, and the counts are reported per dataset (Supplementary Table 8).

### RF raw versus RF+stability ablation and matched control

RF+stability and RF raw were compared at K = 50, 100, 250, 500 and 1,000 using the STRING-only reference for both, in all seven evaluable datasets. Cells in which either network had fewer pairs than K were excluded (PXD005942 at K = 500 and 1,000, where RF+stability has 325 pairs). The RF raw arms of CPTAC_BRCA and PXD005942 were refitted after the configuration defect described above, and their settings and Precision@K values were re-derived with independently written code. To test whether the difference in candidate-pool depth (500 versus 10 predictors per target) explains the gap, the RF raw edge list was truncated per target to its top 10 predictors by importance and re-scored against the same reference, without new inference. The two arms differ in computational effort (B × trees versus trees per target), which this design does not control.

### Prospective replication design

The hypothesis under test was that RF+stability is not necessarily leading at low K but rises to rank 1 by K ≤ 500 in the initial panel. It was recorded before the prospective datasets were selected. Outcome categories, namely full replication (rank 1 at or before the initial panel’s depth threshold), partial replication (qualitatively similar rise, but later) and non-evaluable (not scientifically evaluable under the frozen protocol, independent of any result), were written down after the prospective results had been generated (time-stamped project history), so they are not pre-specified. Because the retrospective pattern itself weakened after the edge-representation correction, the three-way categories do not describe it cleanly, and we report the actual classifications (a qualified replication and a non-replication) instead of forcing the original categories onto the data.

### Bootstrap sensitivity analysis

Because per-iteration edge lists of the primary procedure are not persisted, a sensitivity analysis with a narrower estimator was run on PXD059074 and NAFLD_SomaScan. Per-draw runtime was first estimated from a five-resample pilot (mean 1,147 s per draw for PXD059074; 2,447 s for NAFLD_SomaScan), and B = 100 was chosen as the study’s pre-specified minimum. For each of 100 resamples (seed = 1000 + resample index), samples were resampled with replacement and all five methods were fitted once on the identical resample; random forest was a single untuned fit with edges collapsed to one row per pair, architecturally identical to RF raw. Precision@K (pooled reference) was computed at K = 50, 100, 500 and 1,000. Intervals are 2.5th–97.5th percentiles. Paired differences between random forest and each other method were computed per resample, with two-sided finite-sample p = (n_extreme + 1)/(B + 1), Holm-corrected [20] across the four depths within each dataset and method-pair family; with B = 100 the smallest attainable adjusted value is 0.040. Runs took 6.0 h (PXD059074) and 10.5 h (NAFLD_SomaScan).

### Statistics, software and reproducibility

The depth and rank-reversal analyses are descriptive and no hypothesis test was applied to them; the only inferential statistics are the bootstrap intervals and Holm-corrected paired tests above, which apply to the single-fit estimator. Analyses used R 4.3.3 with ranger 0.18.0, Bioconductor GENIE3 1.24.0, WGCNA 1.74, org.Hs.eg.db 3.18.0 and AnnotationDbi 1.64.1 (versions of the analysis environment; no lockfile was recorded at run time). The pipeline is implemented in R and orchestrated with Nextflow [38] (DSL2, v25.10.4), version-controlled and covered by an automated regression suite (254 expectations, no failures on 29 October 2026). Prospective input matrices were re-verified against the SHA-256 manifest at several points with no discrepancy. Seeds were fixed and recorded for calibration runs and the bootstrap analysis (1000 + index); GENIE3 (all datasets) and the bootstrap stage of RF+stability (five datasets) were not run under a controlled global seed. Implementation defects found and corrected during the work included an input-path assumption affecting two prospective datasets; SOMAmer identifiers reaching enrichment analysis before mapping; a bootstrap driver that read a pre-feature-selection matrix, caught by a dimension check; the zero-as-missing gap; a directed-edge scoring that let RF+stability’s two directions occupy two ranked positions (Supplementary Table 15); and a configuration defect that fitted the RF raw arm of two datasets with one candidate predictor per split. All affected results were regenerated; the corrections are recorded in the code repository.

## Data availability

All datasets are public. CPTAC_BRCA, CPTAC_LUAD, CPTAC_OV and CPTAC_COAD gene-level matrices were obtained from LinkedOmics (linkedomics.org). PXD005942, PXD018996 (originally medRxiv 2020.12.01.20242198) and PXD059074 (source publication: ref. 38) are available from PRIDE/ProteomeXchange under their accessions. NAFLD_SomaScan data are supplementary data to ref. [32]; companion transcriptomic data are at Gene Expression Omnibus (GEO) accession GSE135251. Benchmark tables are provided as Supplementary Tables and the source data for every data figure (Figures 2–5) as Supplementary Data. Canonical and preprocessed matrices, inferred edge lists, checksums and the freeze manifest will be deposited in a public archive before journal publication.

## Code availability

The complete analysis pipeline, including Nextflow workflows, R scripts for data preprocessing, network inference, benchmarking, statistical analyses, and figure generation, is currently maintained in a private GitHub repository. The code will be made publicly available upon publication of the manuscript in journal.

## Acknowledgements

The authors acknowledge the Clinical Proteomic Tumor Analysis Consortium (CPTAC), the PRIDE/ProteomeXchange repository, and the researchers who generated and publicly shared the proteomic datasets analysed in this study. We also thank the developers and maintainers of LinkedOmics, STRING, BioGRID, CORUM, and the open-source software packages used for data preprocessing, network inference, statistical analysis, and benchmarking.

## Author contributions

S.A. (Shahram Aliyari): supervision, conceptualisation, methodology, project administration, writing – review and editing. N.B. (Nazanin Bagherlou): Conceptualisation, methodology, software, formal analysis, data curation, visualization, and writing – original draft. Z.S. (Zahra Salehi): resources, data curation, writing – review and editing.

## Abbreviations

BioGRID: Biological General Repository for Interaction Datasets
BRCA: Breast Invasive Carcinoma
CLR: Context Likelihood of Relatedness
COAD: Colon Adenocarcinoma
CORUM: Comprehensive Resource of Mammalian Protein Complexes
CPTAC: Clinical Proteomic Tumor Analysis Consortium
DIA: Data-Independent Acquisition
GENIE3: Gene Network Inference with Ensemble of Trees
K_common_max: Maximum Common-Support Depth
LFQ: Label-Free Quantification
LUAD: Lung Adenocarcinoma
NAFLD: Non-Alcoholic Fatty Liver Disease
NCI-60: National Cancer Institute 60 Human Tumor Cell Line Panel
OV: Ovarian Cancer
PCA: Principal Component Analysis
PRIDE: PRoteomics IDEntifications Database
RF+stability: Random Forest with Bootstrap Selection-Frequency Filtering
SHA-256: Secure Hash Algorithm 256
SOMAmer: Slow Off-rate Modified Aptamer
STRING: Search Tool for the Retrieval of Interacting Genes/Proteins
TIGRESS: Trustful Inference of Gene REgulation using Stability Selection
TMT: Tandem Mass Tag
WGCNA: Weighted Gene Co-expression Network Analysis

