## Supplementary Information for "Evaluation depth reshapes method rankings in proteomic co-abundance network inference"

All supplementary tables are sheets of Supplementary_Tables.xlsx: - the **README** sheet explains the status labels; - **Column_definitions** defines every column; - **Provenance** gives the source file and SHA-256 of each table.

Tables marked *exploratory* are sensitivity analyses, not confirmatory results.

**Terminology.** The manuscript calls the bootstrap statistic of RF+stability the *selection frequency*. Supplementary tables and source files use the column name stability for the same quantity: the fraction of B bootstrap iterations in which an edge was among its target’s 10 highest-importance predictors.

### Abbreviations

| Abbreviation | Meaning |
| --- | --- |
| B | number of bootstrap iterations |
| CLR | context likelihood of relatedness |
| CPTAC | Clinical Proteomic Tumor Analysis Consortium |
| DIA | data-independent acquisition |
| FDR | false-discovery rate |
| GENIE3 | Gene Network Inference with Ensemble of trees (software name) |
| IPI | International Protein Index |
| K | number of top-ranked protein pairs evaluated |
| K_common_max | smallest network size among the five methods in a dataset (largest fair depth) |
| K_eff | evaluable depth, min(K, network size) |
| KNN | k-nearest neighbour |
| LFQ | label-free quantification |
| MI | mutual information |
| NAFLD | non-alcoholic fatty liver disease |
| PCA | principal component analysis |
| Precision@K | fraction of a method’s K top-ranked pairs present in the reference |
| QC | quality control |
| RF | random forest |
| RF raw | single untuned random-forest fit, top 500 predictors per target |
| RF+stability | random forest with bootstrap selection-frequency filtering |
| TMT | tandem mass tag |
| WGCNA | weighted correlation network analysis |
| Δ | RF+stability minus RF raw Precision@K |

### Supplementary Methods

**Random forest with bootstrap selection-frequency filtering (RF+stability).** - **Base learner (RF raw):** per target protein, a ranger regression on all other proteins, with mtry = ⌊√p⌋, minimum node size 5, the same fixed seed for every fit, and the top 500 predictors by importance. - **Filter:** RF+stability refits this base learner on B bootstrap resamples of the samples and records each target’s 10 highest-importance predictors per resample. - An edge’s selection frequency is the fraction of the B resamples in which it was recorded. - Edges with frequency ≥ 0.60 form the network, ranked by frequency and then by mean importance over the resamples in which the edge was selected. - **Settings** (Supplementary Table 14): B = 20 with 300 trees and permutation importance in six datasets; B = 50 with 500 trees and impurity importance in PXD005942. - **Not error control:** the procedure is a ranking heuristic. Bootstrap resampling with replacement and a fixed number of predictors per target do not provide the error control of subsampling-based stability selection. The negative control (Supplementary Table 17) shows that chance associations can reach high selection frequencies. - **Unequal effort:** RF+stability fits B times as many trees per target as RF raw. - **Seeding:** the bootstrap draws were seeded only in CPTAC_BRCA and PXD005942 (incidentally, by a preceding tuning step of the RF raw arm). GENIE3 was not run under a controlled seed.

**Correction of the RF raw arm (2026-10-29).** - **Defect:** a configuration-merging defect caused the tuning step enabled for CPTAC_BRCA and PXD005942 to fit RF raw with one candidate predictor per split. - **Correction:** both RF raw networks were refitted without tuning, using the same base settings as the corresponding RF+stability arm. - **Checks:** - recorded settings were confirmed; - 10 targets were refitted with plain ranger and compared exactly; - all 30 corrected Precision@K cells were recomputed by independently written code. - **Unaffected:** RF+stability networks, which never used tuning.

**Rank-reversal classification.** - For RF+stability and each other method, the order at the smallest fair depth is compared with the order at every larger fair depth. - Depths at which the two methods have identical Precision@K carry no ordering information and are skipped (primary rule). The rule originally used counted a tie as the second method ahead; both classifications are given in Supplementary Table 2, and one of 24 differs. - The persistent/transient split also depends on where the fair range ends (Supplementary Table 16).

**Common support.** - K_common_max is the smallest native network size among the five methods in a dataset; a depth is fair if K ≤ K_common_max. - In six of seven datasets the limiting network is RF+stability. Its size depends on the selection-frequency threshold, which therefore sets the fair range (Supplementary Table 16).

**Negative control (exploratory).** - **Design:** PXD005942’s preprocessed matrix was permuted protein-wise (20 permutations, seeds 20261101–20261120). The unmodified RF+stability procedure was rerun with the bootstrap seed fixed at 123, and counts were verified by independent code. - **What it removes:** the null removes all associations, including sample-level structure and imputation-induced correlation. Its results are therefore not a false-discovery rate for the observed networks.

**Spearman scoring.** Pairs were ranked by the signed coefficient, so strong negative correlations were ranked last. This convention was used for all reported results.

**Identifier mapping.** Gene-symbol mapping was applied only at the validation boundary. Unresolved identifiers (UNMAPPED_ placeholders and International Protein Index (IPI) accessions) were retained and cannot match the reference (Supplementary Table 8).

### Supplementary Tables

**Supplementary Table 1.** Precision@K, hits, evaluable depth, rank and fair/capped status for 7 datasets × 5 methods × 7 depths, pooled reference. **Supplementary Table 11** gives the full common-support audit; **11b** gives the native network sizes and K_common_max.

**Supplementary Table 2.** Rank-reversal classification of RF+stability against each other method in the six common-support datasets: primary rule (ties carry no ordering; adopted after the results were available) and the rule originally used, with the depths at which ties occur. - For each reversal: the largest Precision@K difference at depths showing the reversed order and at depths showing the initial order. In 7 of 18 reversals the reversed order never exceeds 0.02. **2b:** classes under the superseded directed-edge scoring (historical).

**Supplementary Table 3.** Descriptive decomposition of RF+stability rank changes; not a causal test.

**Supplementary Table 4.** RF+stability versus RF raw (top 500 per target) and the top-10 matched control, STRING reference, K = 50, 100, 250, 500 and 1,000. - Comparisons without equal depth (PXD005942 at K ≥ 500) are excluded. - The superseded defective RF raw values are shown for transparency. - **4b:** summary over equal-depth cells.

**Supplementary Table 5.** Bootstrap sensitivity analysis (B = 100, single untuned random-forest fit per resample, prospective datasets): Precision@K percentile intervals (**5**) and paired differences with finite-sample and Holm-adjusted p-values (**5b**). The smallest attainable adjusted p-value is 0.040.

**Supplementary Table 6.** Preprocessing lineage and parameters for each evaluable dataset, from the recorded quality-control (QC) summaries.

**Supplementary Table 7.** Precision@K against STRING, BioGRID and CORUM separately and pooled (**7**), and method ranks by reference (**7b**). - **7c:** chance-level precision per dataset, i.e. the fraction of all pairs of measured proteins present in the pooled reference (0.9–1.5% in the six cross-method datasets, 6.7% in PXD005942).

**Supplementary Table 8.** Identifier coverage: analysis-matrix features resolved to gene symbols (95.2–100%) and RF+stability pairs retained for scoring.

**Supplementary Table 9.** Recorded wall-clock runtimes (indicative; different runs, machines and loads), including the corrected RF raw single fits.

**Supplementary Table 10.** PXD018996 evaluability-gate metrics recorded before any inference (2026-09-07).

**Supplementary Table 12.** Missingness and evaluability boundary for all eight datasets. The PXD018996 sample counts are reproduced as recorded; the table notes their inconsistency.

**Supplementary Table 13.** RF+stability network size in *directed* edges at selection-frequency thresholds 0.3–0.8 (six datasets) (**13**); distribution of selection frequencies among retained edges (**13b**); PXD059074 Precision@K with 300 versus 240 trees (**13c**).

**Supplementary Table 14.** Per-dataset configuration as executed: trees, mtry, importance type, B, threshold, seeding.

**Supplementary Table 15.** Superseded directed-edge scoring versus the corrected collapsed scoring for RF+stability (historical comparison).

**Supplementary Table 16 (exploratory).** Stricter selection-frequency thresholds (0.7, 0.8, 0.9, 0.96) applied to the stored RF+stability networks without refitting. - **Reported:** network sizes, K_common_max, fair depths, leaders and reversal classes. - **Finding:** no Precision@K value at a fair depth changes; only the fair range does. - 0.6 remains the primary analysis, and no threshold was selected from these results.

**Supplementary Table 17 (exploratory).** PXD005942 negative control. - **17:** directed and undirected edge counts passing each threshold in 20 complete-null permutations versus the observed network. - **17b:** pooled-reference Precision@K of the null networks. - Not an estimate of false-discovery rate.
